# How young is your Muscle? A Machine Learning framework for motor functional assessment with ageing by NMF based analysis of HD-sEMG signal

**DOI:** 10.1101/2020.02.12.946343

**Authors:** Swati Banerjee, Sofiane Boudaoud, Kiyoka Kinugawa-Burron

## Abstract

**Objective:** With ageing, there are various changes in the autonomic nervous system and a simultaneous decline in the motor functional abilities of the human body. This study falls within the framework improvement of the clinical tools dedicated to the robust evaluation of motor function efficiency with ageing.

**Method:** Analysis of HD-sEMG signals recorded from 32 channels during Sit To Stand (STS) test are used for the functional assessment of body muscles. For this purpose, five primary characteristic features, *iEMG, ARV, RMS, Skewness, Kurtosis*, are employed for the study. A channel clustering approach is proposed based on the parameters using Non Negative Matrix Factorization (NMF).

**Results:** The NMF based clustering of the HD-sEMG channels seems to be sensitive toward modifications of the muscle activation strategy with ageing during STS test.

**Conclusion:** This manuscript provides a framework for the assessment of Motor Functional Age(MFA) of subjects having a range of chronological from 25 yrs to 75 yrs. The groups were made a decade apart and it was found that the MFA varies with the level of activeness of the muscle under study and a premature ageing is observed according to the change in activation pattern of the HD-sEMG grid.

## 1. Introduction

According to the World Health Organization (WHO)’s report on the Global health and Ageing wor (2015), the geriatric population will have a sudden inflation in the next few decades, demanding prompt healthcare facilities. To be efficient, these amenities need a precise screening of musculoskeletal system. Also, due to sedentary lifestyle and inactivity the younger generation is suffering from various threatening health disorders. One such problem is premature muscle ageing and this manuscript focuses on addressing it by proposing a probable solution to screen the muscular efficiency of an individual. The musculoskeletal system is a complex arrangement of muscles and other organs that accounts for 40% of body mass. Human body has more than 600 muscles of varying size and shape Basmajian (1962) providing support to the body and enabling movement abilities. Ageing is associated with a large number of physiological Sharples A. P. (2018) and neuromuscular Booth et al. (1994) changes. These phenomenon can be observed by screening the functional capacity of muscles which usually tend to decreases with ageing. Surface Electromyography (EMG) refers to the collective electrical activity recorded, non-invasively from the muscles, produced during muscle contraction by the motor units (MU) and is being controlled by the Central Nervous System (CNS) through the peripheral neural system (PNS). This signal corresponds to the summation of MU’s action potential trains within the recording range of the electrode. In this research, we use the High Density sEMG (HD-sEMG) technique which is a spatio-temporal variant of the monopolar single channel or bipolar sEMG signalsNaik et al. (2014),Arjunan et al. (2015), Van Dijk Johannes et al. (2012).

Milner-Brown and Stein found that the Average Rectified Value (ARV) of the MUAP varies approximately linearly with the corresponding force Milner-Brown and Stein (1975) and Moritani and deVries Moritani et al. (1978) confirmed the linear theory by exploring the relationship between the integrated EMG (iEMG) and the net joint force. M. Al. Harrach et al. Al Harrach et al. (2017) has used both higher order statistics and robust functional statistical features on the probability density function shape screening of the sEMG data.

This manuscript explores the ability of HD-sEMG technique to study the behaviour of the rectus femoris muscle as a function of ageing categorization using a Sit to Stand (STS) test. Ten primary characteristic features, *iEMG, ARV*, *RMS*, *Skewness, Kurtosis*, values are estimated for the study. These features are significant enough to discriminate among the ageing categories under consideration Al Harrach et al. (2017). A non negative matrix factorization based channel selection approach is used to identify the muscle contraction strategy in the rectus femoris muscle in the three ageing categories. This method was also explored by Gazzoni et al. (2014) for identifying forearm muscle activity during wrist and finger movement.

## 2. Experimental protocol and Data acquisition

### 2.1. Experimental Protocol

The aim of this experiment is to design a STS protocol to realise the study of chair lift movement on the right Rectus Femoris muscle contraction. The motivation behind using this particular muscle is two fold, firstly because of its anatomical position on the lower part of the upper limb, secondly because of the fact that it is the most solitary muscle involved during the daily postural changes in the lower limbs Booth et al. (1994) and also because this muscle gets activated during the STS test over a chair. All the participants were made aware of the protocol and their consent were taken before recording the HD-sEMG. A standard sized chair with an approximate seat height of 45cm and seat size of 40×40cm was used for the experiment. Data was recorded in a room with a comfortable normal temperature. Figure 1. shows the protocol in action for the acquisition of the HD-sEMG dataset.

**Figure 1:**
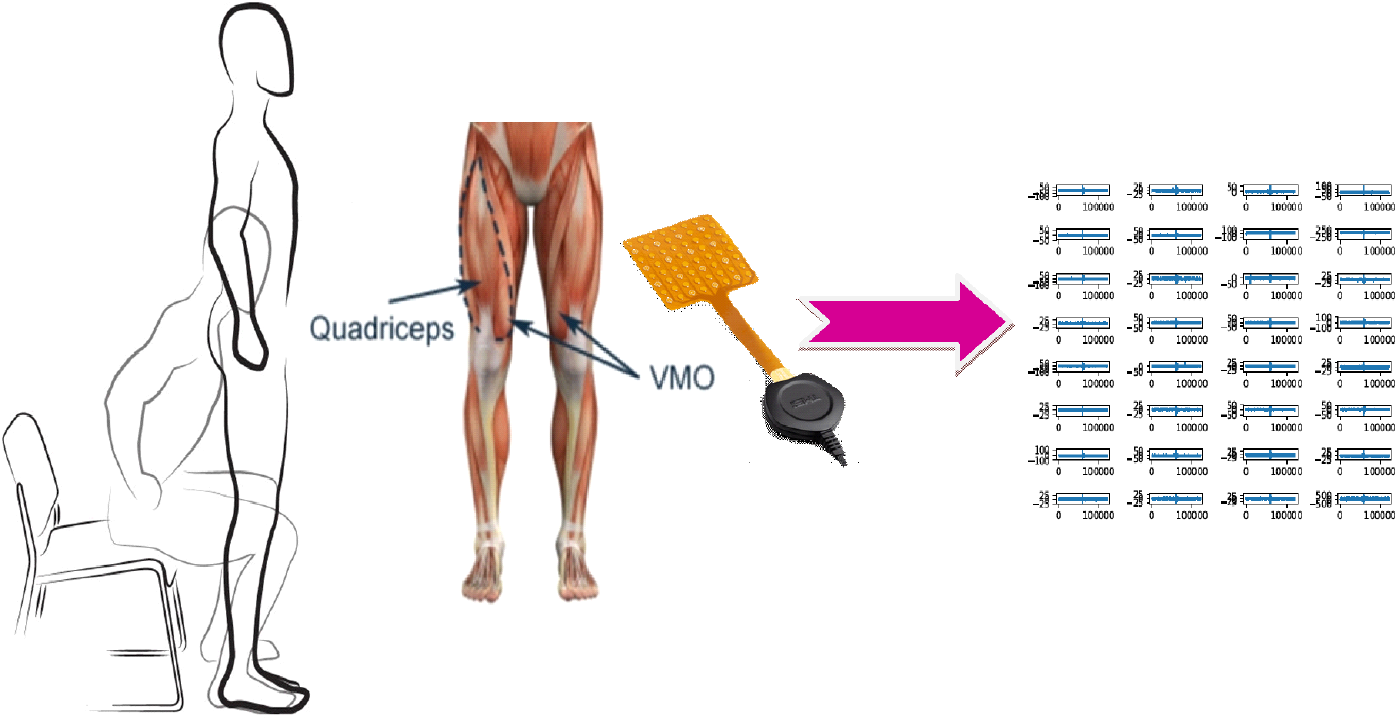
Sit to Stand Chair Test

### 2.2. Participants

In this study, 82 healthy participants were recruited to perform a Sit-To-Stand(STS) test. The participants did not have any history of muscular or neuro-physiological disorder and were known to have lower limbs with medically normal functionalities. To asses the physical activity level of the participants they have to compile International Physical Activity Questionnaire (IPAQ) mentioning about their daily routine and lifestyle during the precedent 7 days. IPAQ is a medically recognised instrument designed mainly for population surveillance of physical activity among adults Craig et al. (2003); Bassett Jr (2003). Scores based on the walking, moderate-intensity and vigorous intensity activity were computed and subjects having a certain score was accepted to be included in the cohort study. All volunteers gave their free and informed consent for the experiments and were able to complete the protocol accordingly. Table 1 shows the available group of subjects for this study. Subjects with a *BMI >* 26.5 were not included in the study.

**Table 1:** Grouping of Participants

| No. of Participants | Age(years) |
| --- | --- |
| 15 (9M, 6F) | $27.133 \pm 2.799$ |
| 19 (7M, 12F) | $40.052 \pm 3.964$ |
| 16 (7M, 11F) | $50.689 \pm 4.346$ |
| 19 (10M, 8F) | $60.684 \pm 3.637$ |
| 13 (7M, 6F) | $69.307 \pm 1.750$ |

### 2.3. Data Acquisition

Three sets of STS(Sit To Stand) tests were performed for each of the participants at a spontaneous pace. Figure 1 shows the HD-sEMG data acquisition scheme. All subjects were able to perform and complete the protocol with very little training and explanation.

#### 2.3.1. High Density Surface EMG (HD-sEMG)

The acquisition is performed through a 32-electrode square grids (4×8), fixed in a relevant manner aligned to the muscle of interest for our case the rectus femoris muscle, using a multichannel amplifier (32 acquisition Channels) called MOBITA designed by TMSi© which is characterized by 4 mm diameter electrodes and an inter-electrode distance of 8.57 mm.

#### 2.3.2. Acceleration Data to monitor the STS protocol

To monitor the movement cycle, along with the HD-sEMG data, acceleration data was also acquired simultaneously. It was found that maximum acceleration in Y direction was of significance for this analysis.

## 3. Method

The flowchart for the Machine Learning framework used in this work is depicted in Figure 3. It comprises of three primary blocks, explained as: a. The Signal Processing and Feature extraction: The acquired 32 channel HD-sEMG signals were pre filtered and segmented. Features as mentioned in section 3.3 were extracted along with some frequency domain features like, Average Power Spectral Density (APSD), Mean Frequency (Mean Frequency), Energy of the signal etc. Some other features like the Zero Crossing(Z-Crossing) and signal duration was also computed. A total of 10 Features were extracted from the database. b. Learning by Clustering: Once the Feature matrix are fixed, they were subjected to Unsupervised NMF which clusters the HD-sEMG channels. To understand the channel configuration four parameters are extracted from the Grid as stated in the next section. Accelerometer Data (Max Accel) was fused along with the Sit To Stand Duration (STS) duration to feed into the next classification block. c. Classifier Testing and Training: Once the Grid Score are established six classifiers are used to identify the class of the subjects. In this work, the data was acquired during the STS protocol as explained in sec 2.3. During the preprocessing phase the signals were visually inspected and proper care has been taken to make the signal as artefact free as possible. The time domain and the statistical features were estimated using the equation(1)–(6). Along with these features the STS was recorded. MOBITA allows the recording of maximum acceleration and this measurement was also preserved. Once all the features are extracted NMF based channel clustering was performed.

**Figure 2:**
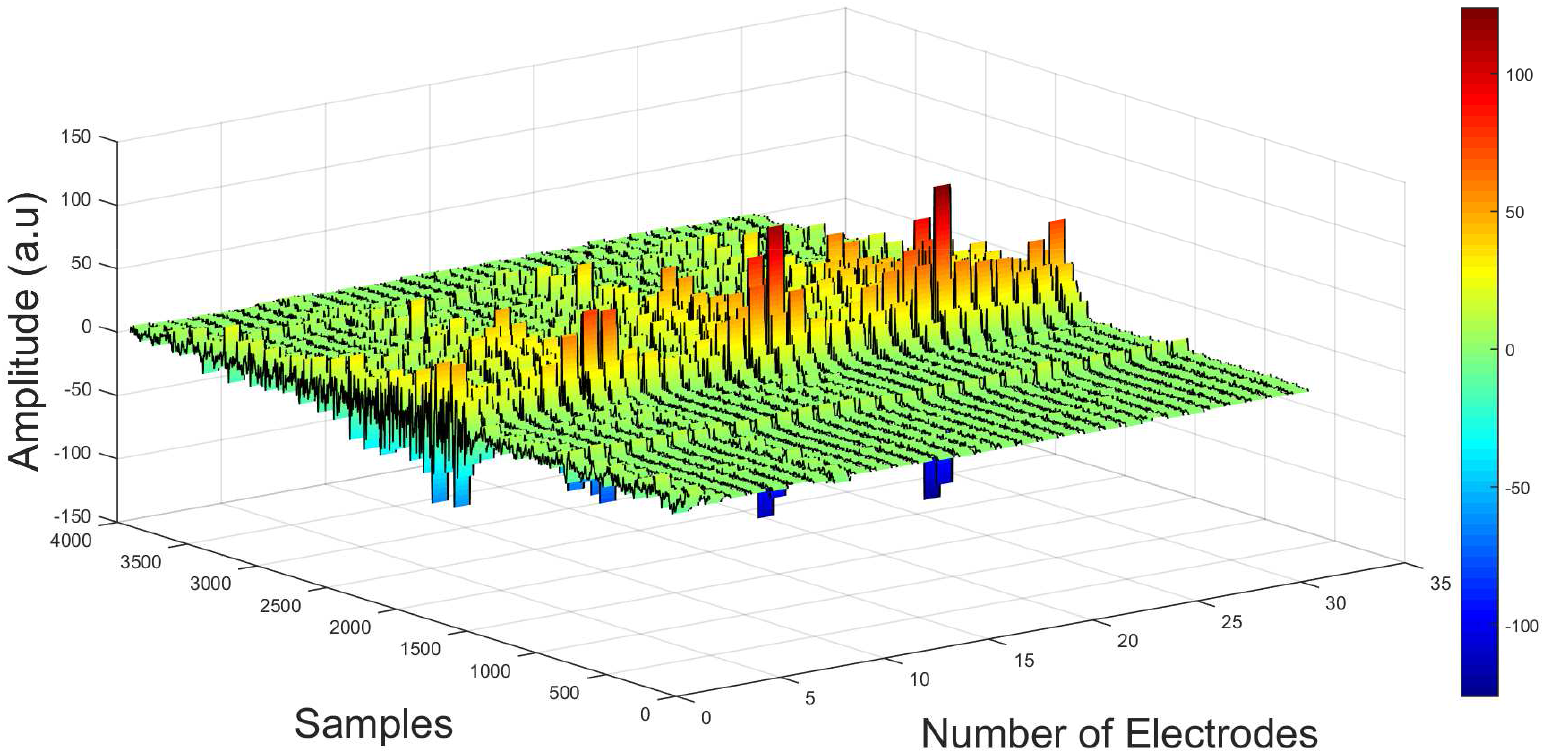
Spatial View of a 32 channel HD-sEMG data

**Figure 3:**
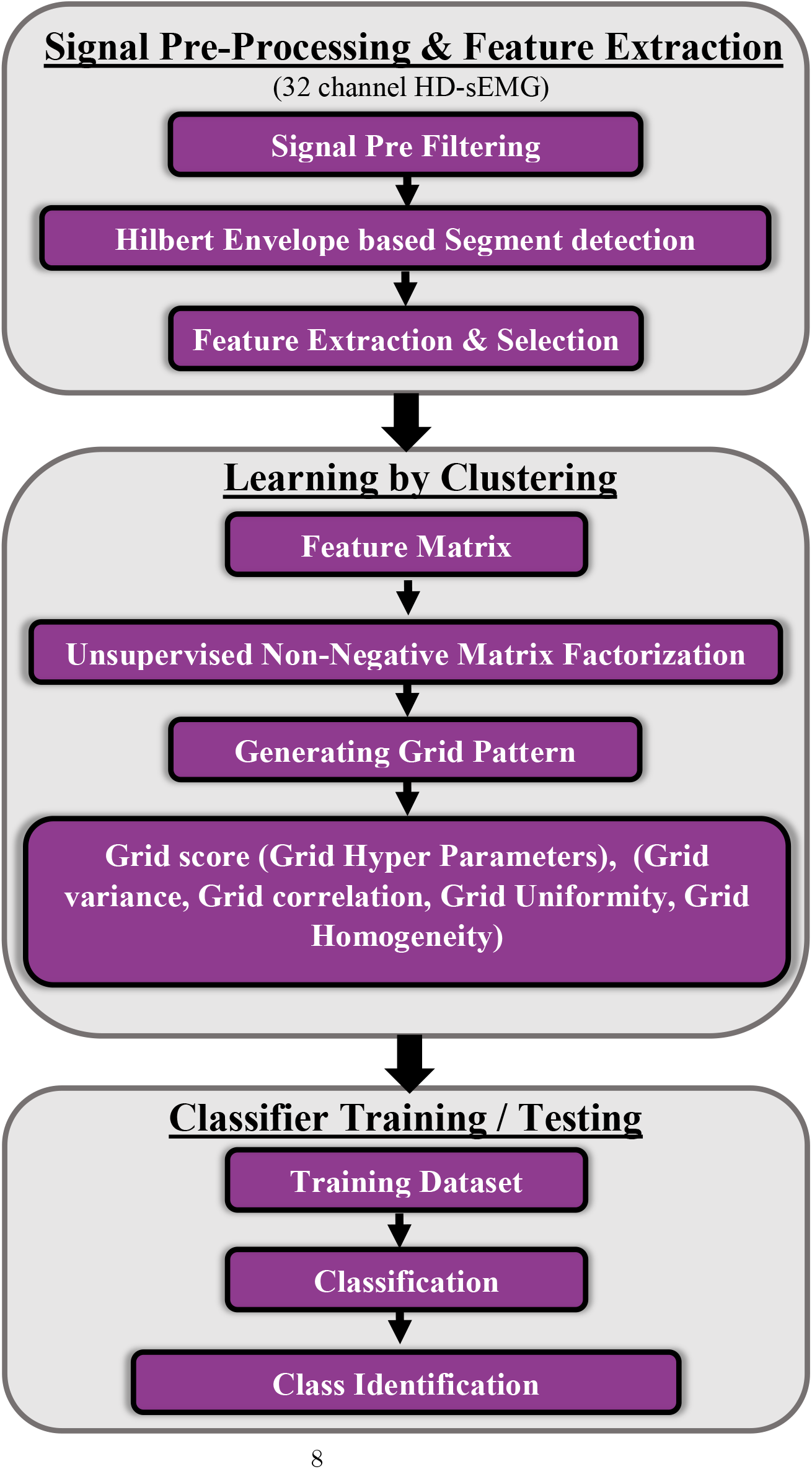
A schematic View of the proposed Machine Learning Framework

### 3.0.1. Pre Filtering and Hilbert based segmentation of HD-sEMG Contraction

The HD-sEMG signals were digitally filtered with a 4^th^ order Butterworth filter with a cut-off range of 20-450 Hz and powerline interference(50Hz and its four higher harmonics) was removed using spectral interpolation (Mewett et al. (2004)). An adaptive threshold based on the hilbert envelope of the HD-sEMG signal was used to extract the dynamic contraction segment from the EMG data extracted from each channel. This is to avoid the influence of noise that may occur in the remaining signal other than the contraction segment and also to get an exact duration of the contraction and hence the explore the muscle activation strategy.

### 3.0.2. Feature Analysis and selection

The time domain and statistical features extracted from the 32 channel HD-sEMG data for quantification of the muscle activation strategy of the Rectus Femoris muscle are described below, where all the symbols have there usual meaning. These features were selected taking into account their nature of revealing relevant information from the signal under consideration (?) and are known to be prominent parameters related to HD-sEMG related studies (?).

*ARV*: The average rectified value (ARV) gives the average of the rectified values of a signal *s_i_* during a segment of time corresponding to the samples, say *N_tot_*,

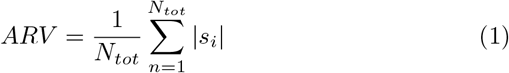
*RMS*: The RMS is a popular amplitude/time domain feature in the EMG signal analysis also known as ‘EMG amplitude detector’. This popularity is due it’s capacity to assess the main force generated by the muscle through estimation of the level of contraction [197][199]. It is an indicator of the health situation of the patients and is known to give a difference between healthy individual and a person having some muscular dysfunction as it correlates with the power of given EMG signal:

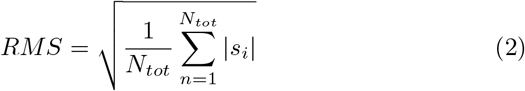
*Skewness*: Skewness value was computed using,

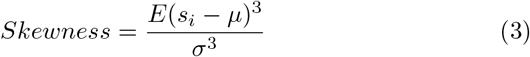
*Kurtosis:* Kurtosis value was computed using,

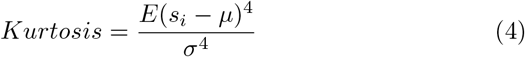
*iEMG:* The integrated EMG which is the integral of a digitized rectified signal *s_i_* for a period of time corresponding to the total number of samples given by *N_tot_*. It is the indication of measure of muscle activity or effort during a given duration of the signal. More amplitude, duration and frequency of action potentials lead to the large value of integration. It is the best measure for the total muscular effort and is given by:

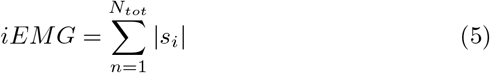
*APSD*: Average Power Spectral Density. The power spectrum of an EMG signal describes the distribution of the signal’s power with frequency and is calculated by squaring the Fourier Transform of each segment of data and averaging them. This gives a measure of the power that each frequency contributes to the EMG signal.For our analysis Mean Power of the signal is computed.

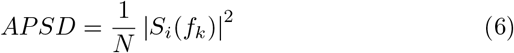

where, |*S_i_*(*f_k_*)| is the *N* point Fourier transform of the signal *s_i_* acquired by the *i^th^* electrode.
*SigEn:* Energy of the signal gives an estimation of the strength.

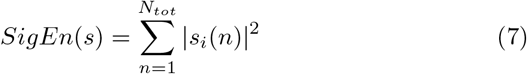

where n= number of samples.
*SigDu:* Signal duration was computed by measuring the duration of the contraction segment as extracted from the hilbert envelope method explained in section 3.1.1.
*Z-cross:* Number of zero crossings is a feature counting number of times a signal crosses the axis of abscissas (zero axis). The random temporal fluctuations of the sEMG signal may serve as distinguishable feature performing the detection of diseases and is given as:

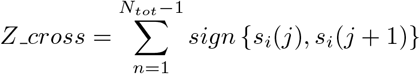

where,

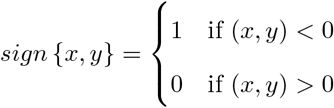 All these features were computed on a specific time window during the STS exercise, with all the notations having usual meaning.

### 3.1. Learning By Clustering and Data fusion From other Modalaties

So we have a feature matrix of *F* = 31 × 9 for each trial set of HD-sEMG data set. One of the 32 channel is used as the electrical reference and hence not considered for analysis.

#### 3.1.1. Non Negative Matrix Factorization based Channel Selection

Non-negative matrix factorization (NMF) has been successfully applied in the mining of biological data Brunet et al. (2004); Lee and Seung (1999). This manuscript aims to address the issue of characterization of the ageing categories of the five available classes as in table 1. To cater this requirement, an EMG channel reduction approach is developed using NMF based channel clustering.

NMF is a matrix decomposition approach which decomposes a non-negative matrix *X* ∈ *R*^*m×n*^ into two low-rank non-negative factors *A* ∈ *R*^*m×k*^ and *Y* ∈ *R*^*m×k*^ where (*k* < *min*(*m,n*)), that is

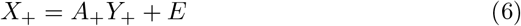

where, *E* is the error or the residual and *M*_+_ indicates the matrix is non negative, and its optimization in the euclidian space is formulated as:

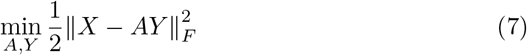

subject to, *A, Y* Statistically, this formulation is obtained from the log-likelyhood function under the assumption of a finite Gaussian error. Assuming that the multivariate data points are arranged in the columns of *X*, then *A* is called the *basis matrix* and *Y* is called the *Coefficient matrix*;each column of *A* is thus a *basis vector*.

##### NMF based Channel Clustering

NMF has been know to have clustering properties and in this manuscript we have explored this property for selection of the channels most relevant to the study. The selection of the channel can reveal many patho physiological phenomenon related to the activation of the rectus femoris in case of three categories of participants. Given the data *X* with multivariate data points in the columns, the idea is that after applying NMF on *X*, a multivariate data point, say *x_i_* is a non-negative linear combination of the columns of *A*; that is

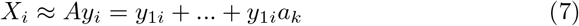

The largest coefficient in the *i^th^* column of *Y* indicates the cluster that this data point belongs to and this is due to the fact that the data points are mainly composed of the main basis vectors and hence they should therefore be in the same group. A basis vector is usually looked upon as a cluster centroid. For this particular study the clustering of the HD-sEMG channels gives an estimate of the activation of the region for rectus femoris muscle. The number of clusters can give an insight into the modification of the muscle activation strategy and their clinical significance in case of ageing categories.To preserve the spatial information of the electrode clusters they were arranged in the 8 × 4 grid shape to resemble the original pattern of the acquisition Electrode array arrangement.

### 3.2. Grid Score Computing and Data Fusion

#### 3.2.1. Electrode Grid Score Estimation

Parameters estimated based on the clustering of HD-sEMG channels using semi-NMF. The clustered electrodes can be visualised as a Grid image, say *G*. To further understand the spatial relationship among the 8 × 4 HD-sEMG electrodes placed over the rectus Femoris muscle, four statistical features are estimated. The value for *e*32 = *g*(1, 4) = 0, as it is the reference electrode used during EMG data acquisition. This grid image can be a probable representation of the collective motor recruitment and excitation pattern.

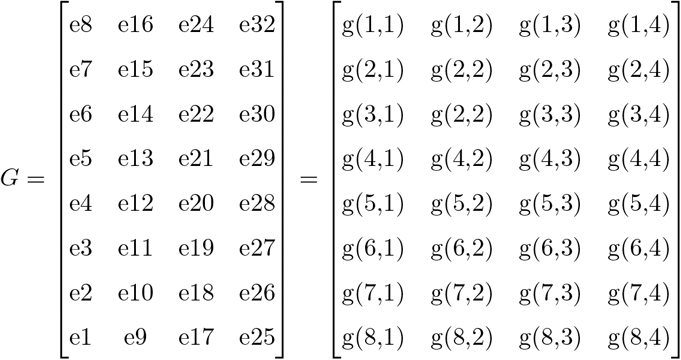

The features are enlisted in eq(8)–eq(11). Here (*i,j*) represents a particular pixel element.

- Variance (GV): Also known as the grid contrast is the measure of the intensity contrast between a pixel and it’s neighbour over the whole image (in this case the electrode grid).

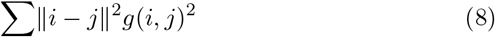
- Correlation (GC): It gives the statistical measure of how correlated a pixel is to it’s neighbour over the whole image range. Correlation is 1 or −1 for a perfectly positive or negatively correlated image. Correlation is NAN for a constant image.

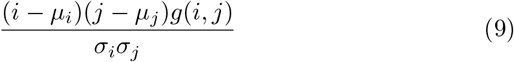
- Uniformity (GU): It is the representation of the energy content of the grid image. Energy is 1 for a constant image.

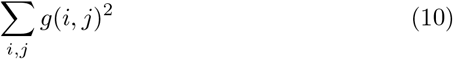
- Homogenity (GH): It is the closeness of distribution of the elements in the grid image and is given by

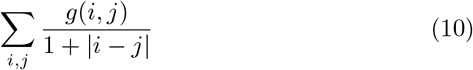

#### 3.2.2. Data fusion from other Significant Measurements

The maximum value of the Trunk acceleration data *MaxAccel* in the Y direction and the Sit to Stand duration, *STS* was found to be discriminative for this protocol A set of hyperparameter is thus generated as in tuple *GS* = {*GV, GC, GU, GH, MaxAccel*}

### 3.3. Classification Techniques and Data Normalization

- Normalization In machine learning and data mining, data processing plays a key role in optimizing the predictive performance.Hence, normalization approach is applied as a data preprocessing before Classification steps. ? explained that the 2 main goals of data pre-processing are data quality optimization, and data normalization.
- MultiClass LogistiC Regression (MLRC) In statistics, multinomial logistic regression is a classification method that generalizes logistic regression to multiclass problems, i.e. with more than two possible discrete outcomes. That is, it is a model that is used to predict the probabilities of the different possible outcomes of a categorically distributed dependent variable, given a set of independent variables (which may be real-valued, binary-valued, categorical-valued, etc.).
- Linear Discriminant Analysis (LDA): Generally, it discovers the most discriminant projection by maximizing between-class distance while minimizing within-class distance. Different class labels can be specified using the membership matrix.
- K-Nearest Neighbor (KNN): It is a non-parametric machine learning method that can be applied for different purposes. This algorithm keeps entire existing cases, while new cases will be classified based on a similarity measure. It applies the construction of a multi-dimensional feature space, where the various dimensions are associated with different signal features (e.g. standard deviation or mean).
- Classification And Regression Tree (CART) CART also called Decision tree learning is a method commonly used in data mining. The goal is to create a model that predicts the value of a target variable based on several input variables.
- Naive Bayes Classifier (NB) It is an extremely practical a probabilistic classifier. It performs learning tasks where each sample *x* is explained by using a conjunction of attribute values. Accordingly, the target function *f* (*x*) can take various values from the finite set.
- Support Vector Machines (SVM) Linear SVM is the newest extremely fast machine learning (data mining) algorithm for solving multiclass classification problems from ultra large data sets.

### 3.4. Performance Evaluation and Statistical Analysis

Once the parameters are extracted they were analysed for statistical significance using ANOVA test, where only (*p value* < 0.05 * and *p value* < 0.001**) are considered as significantly different for identification of the various ageing categories. Table 2 shows the overall results in terms of median and the median absolute deviation (MAD) of the dataset.

**Table 2:**
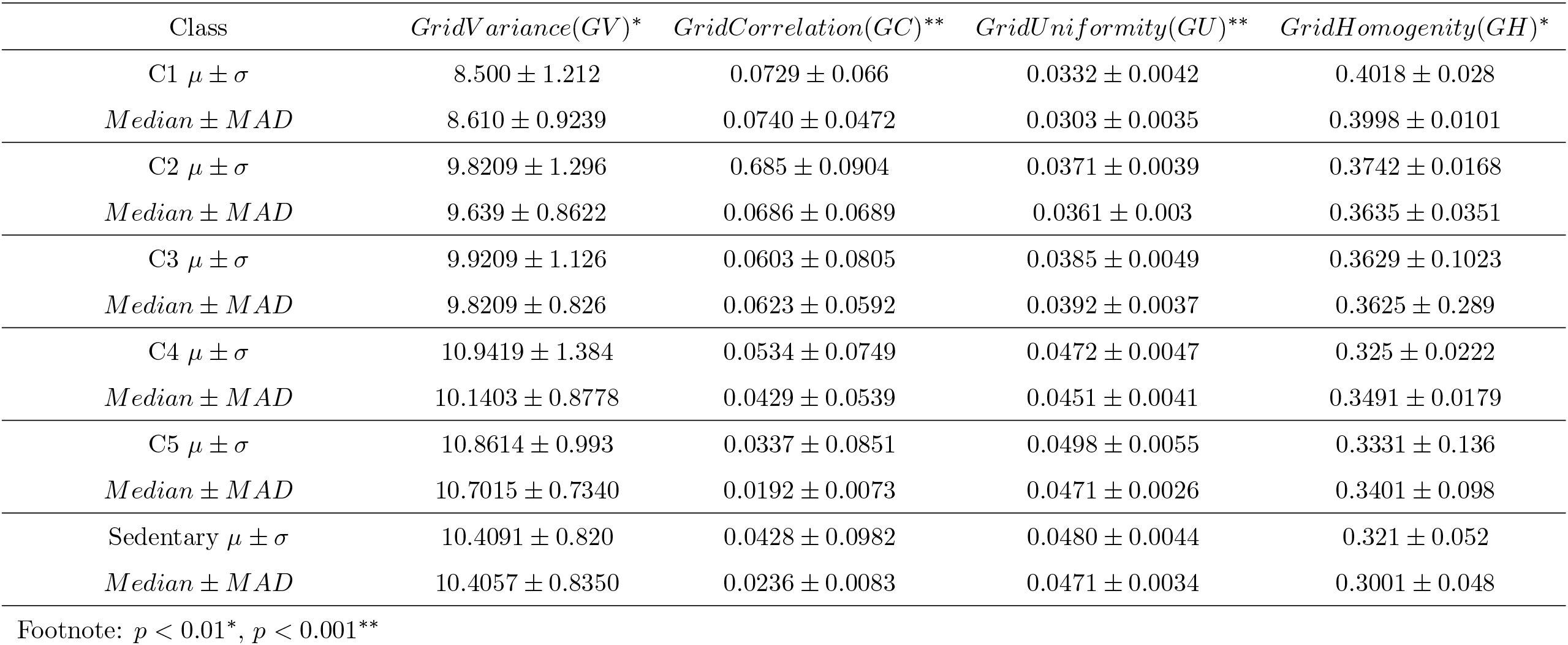
ANOVA Test on the Grid HyperParameters

| Class | $GridVariance(GV)^*$ | $GridCorrelation(GC)^{**}$ | $GridUniformity(GU)^{**}$ | $GridHomogeneity(GH)^*$ |
| --- | --- | --- | --- | --- |
| C1 $\mu \pm \sigma$ | 8.500 ± 1.212 | 0.0729 ± 0.066 | 0.0332 ± 0.0042 | 0.4018 ± 0.028 |
| *Median ± MAD* | 8.610 ± 0.9239 | 0.0740 ± 0.0472 | 0.0303 ± 0.0035 | 0.3998 ± 0.0101 |
| C2 $\mu \pm \sigma$ | 9.8209 ± 1.296 | 0.685 ± 0.0904 | 0.0371 ± 0.0039 | 0.3742 ± 0.0168 |
| *Median ± MAD* | 9.639 ± 0.8622 | 0.0686 ± 0.0689 | 0.0361 ± 0.003 | 0.3635 ± 0.0351 |
| C3 $\mu \pm \sigma$ | 9.9209 ± 1.126 | 0.0603 ± 0.0805 | 0.0385 ± 0.0049 | 0.3629 ± 0.1023 |
| *Median ± MAD* | 9.8209 ± 0.826 | 0.0623 ± 0.0592 | 0.0392 ± 0.0037 | 0.3625 ± 0.289 |
| C4 $\mu \pm \sigma$ | 10.9419 ± 1.384 | 0.0534 ± 0.0749 | 0.0472 ± 0.0047 | 0.325 ± 0.0222 |
| *Median ± MAD* | 10.1403 ± 0.8778 | 0.0429 ± 0.0539 | 0.0451 ± 0.0041 | 0.3491 ± 0.0179 |
| C5 $\mu \pm \sigma$ | 10.8614 ± 0.993 | 0.0337 ± 0.0851 | 0.0498 ± 0.0055 | 0.3331 ± 0.136 |
| *Median ± MAD* | 10.7015 ± 0.7340 | 0.0192 ± 0.0073 | 0.0471 ± 0.0026 | 0.3401 ± 0.098 |
| Sedentary $\mu \pm \sigma$ | 10.4091 ± 0.820 | 0.0428 ± 0.0982 | 0.0480 ± 0.0044 | 0.321 ± 0.052 |
| *Median ± MAD* | 10.4057 ± 0.8350 | 0.0236 ± 0.0083 | 0.0471 ± 0.0034 | 0.3001 ± 0.048 |

Footnote: $p < 0.01^*$ , $p < 0.001^{**}$

### 3.5. Data Classification and Crossvalidation

After the preprocessing, we divide the data into training sets and testing sets based on the stratified 10-fold cross-validation method. The use of k-fold cross validation method has the following advantages: a)reduced the overfitting effect, b) made the results more reliable, c) increased the efficiency of feature selection.

## 4. RESULTS

The Python language along with Pandas, and Sklearn library were used in this work. We used an Intel Core i5-7300HQ 3.5 GHz machine with 16 GB of RAM and single core.

### 4.1. Hilbert Envelope based Contraction Segmentation

**Figure 4:**
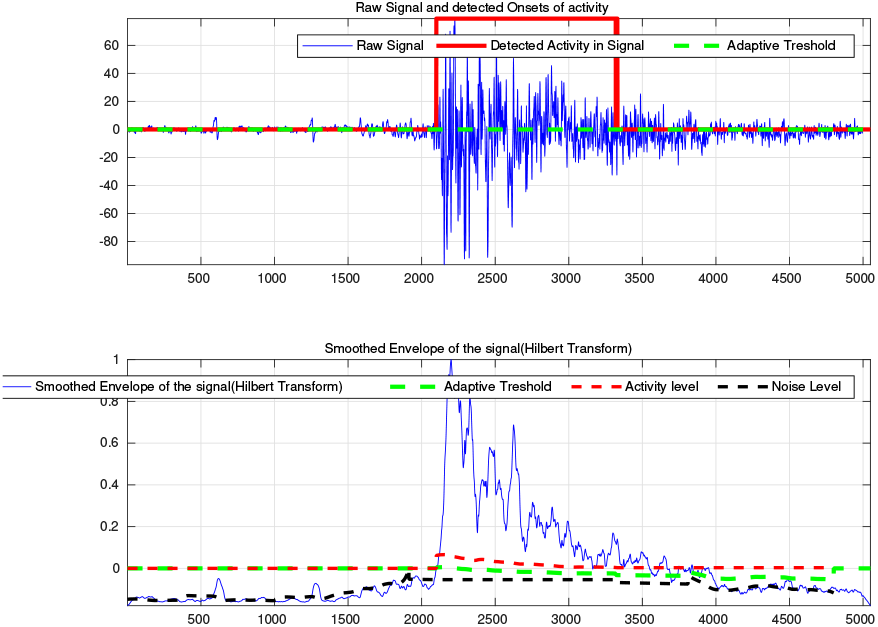
EMG contraction Extraction.

### 4.2. HD-sEMG Signal Preprocessing and Feature Extraction

This part is intentionally Not included in the text but will be available in the Next version.

### 4.3. Analysis of Grid Hyper Parameters

The Grid Hyper-Parameters were computed and were tested for all the Class. Table 2 shows the results of the ANOVA test. It was found that the Correlation and Energy Parameters are the most discriminative among groups. It is evident from Table 3 that the Sedentary Class which actually belongs to the ageing group of C3 has parameter characteristic similar to that of C4 and C5. This can be due to the fact that the grid score as explained in detail in section 2.3, obtained from the analysis explores the HD-sEMG grid homogeneity and with the progression of ageing the in-homogeneity is enhanced giving an indication of the about the prognosis of ageing phenomenon.

**Table 3:**
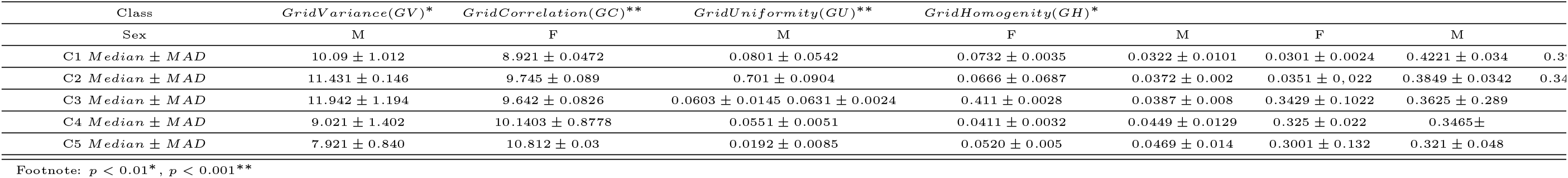
ANOVA Test on the Grid HyperParameters

### 4.4. Classification Results

After the preprocessing, we divide the data into training sets and testing sets based on the stratified 5-fold crossvalidation method. The use of k-fold cross validation method has the following advantage: (a) reduced the overfitting effect, (b) made the results more reliable, (c) increased the efficiency of feature selection and parameter optimization. An average accuracy over all the fold is reported in this manuscript.

## 5. Discussion

The statistical and time domain feature descriptors thus obtained in section 3.1, have a significant sensitivity over the three ageing groups mentioned in table 1. The ANOVA test shows that the characteristic features thus obtained are statistically relevant with a *p* < 0.001 ?. Figure. 2 shows the ANOVA test of the maximum amplitude feature of the three classes of the Dataset as mentioned in table 1. Figure 3, shows the spread of the extracted features each from the three ageing groups. Figure 4, depicts the clustering of 31 Channels marked with values from ‘1 to 5’. A ‘0’ shows channel no. 32, used as the reference during data recording. The results shows that the clusters are comparatively homogeneous for the class 1 and class 2 cases, however, for the ageing group 57 ± 7.07 there are more clusters available, giving a probable indication of the change in muscle activation phenomenon with ageing. The features from each channels of the HD-sEMG signal are representative of underlying homoeostasis of the studied muscle and the clustering of channels explores the inter-channel correlation hence giving an insight into localised activation of rectus femoris muscle during the STS test. Furthermore, this clustering results explores ways of functional assessment of the rectus femoris muscle with ageing. With ageing the muscle might lose its efficiency and strength, likewise there is a deterioration in coordination among various regions of the muscles.

**Figure 5:**
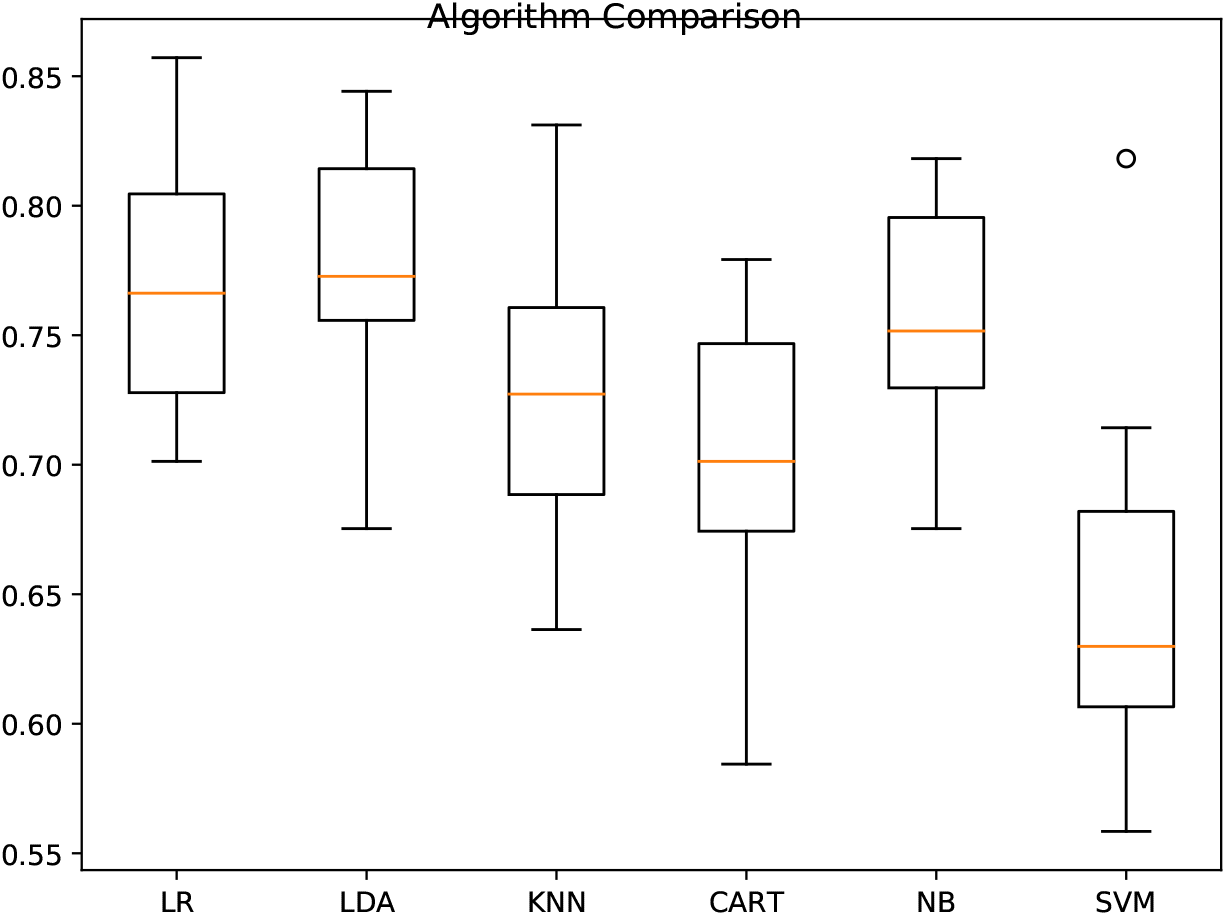
Comparison of Various Classifier accuraccy

